## Supplementary material for "Adult patients with autoinflammation of unknown origin partially phenocopy the immune presentation of Still’s disease"

**Supplementary Spreadsheet 1. Clinical resource for SAID patients.** Clinical and demographic features for each included SAID patient in the cohort.

**Supplementary Spreadsheet 2. Gated populations with gating hierarchy.** Gating hierarchy and marker definitions for the 208 immune variables (frequency of population within parent or super-parent) used in the study.

**Supplementary Spreadsheet 3. Flow cytometry data resource for SAID patients.** Raw flow cytometry cell frequency for immunological subsets across the patients and controls used in this study, together with demographic features.

**Supplementary Figure 1. Cells populations with high association with autoinflammation of unknown**. Odds ratio and 95% confidence interval of 50 highly-associated cell population frequency changes in patients with inflammation of unknown origin in relation to healthy individuals. Estimated by multivariable logistic regression adjusted by sex and age

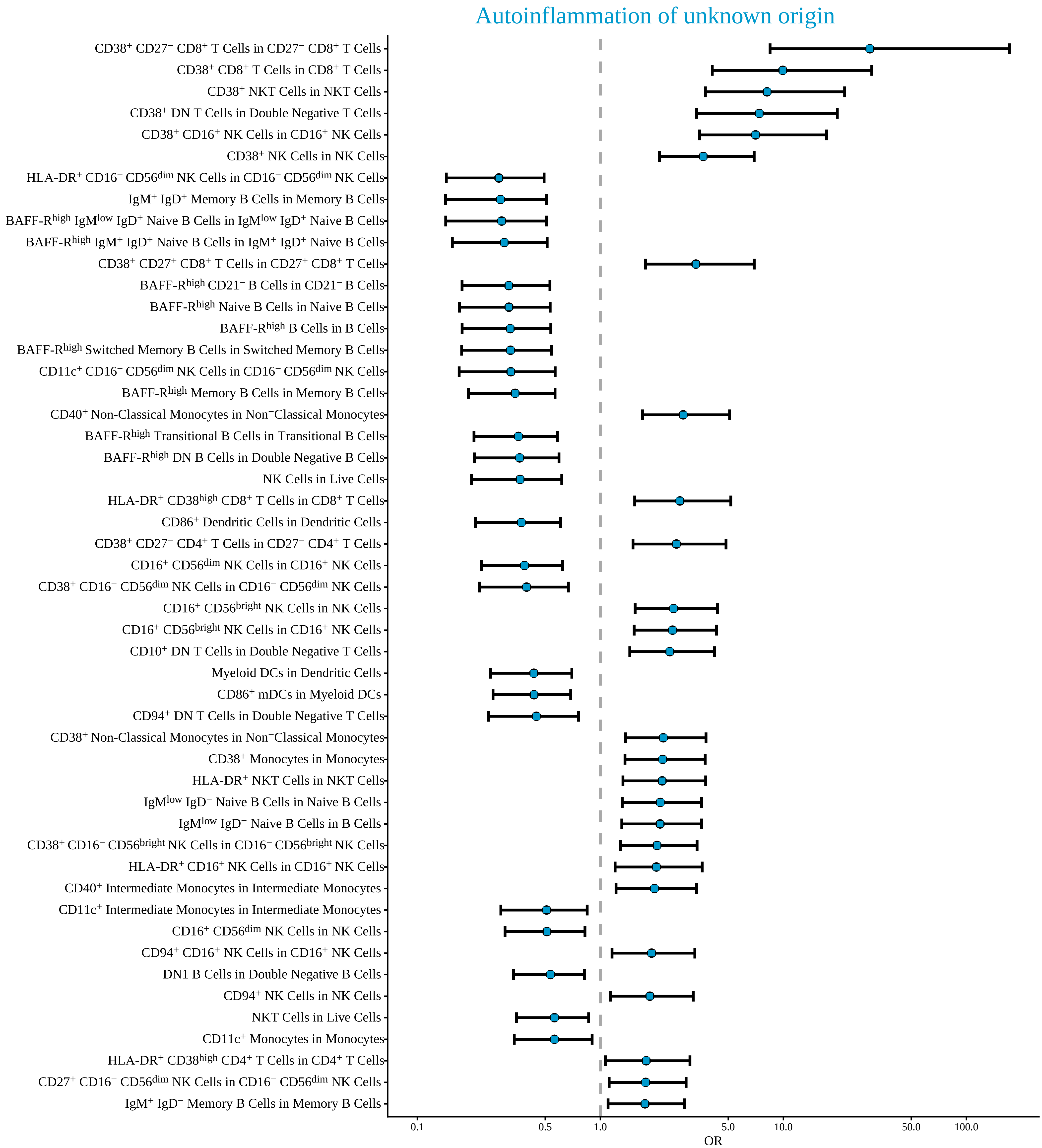

Autoinflammation of unknown origin
CD38+ CD27− CD8+ T Cells in CD27− CD8+ T Cells
CD38+ CD8+ T Cells in CD8+ T Cells
CD38+ NKT Cells in NKT Cells
CD38+ DN T Cells in Double Negative T Cells
CD38+ CD16+ NK Cells in CD16+ NK Cells
CD38+ NK Cells in NK Cells
HLA-DR+ CD16− CD56dim NK Cells in CD16− CD56dim NK Cells
IgM+ IgD+ Memory B Cells in Memory B Cells
BAFF-Rhigh IgMlow IgD+ Naive B Cells in IgMlow IgD+ Naive B Cells
BAFF-Rhigh IgM+ IgD+ Naive B Cells in IgM+ IgD+ Naive B Cells
CD38+ CD27+ CD8+ T Cells in CD27+ CD8+ T Cells
BAFF-Rhigh CD21− B Cells in CD21− B Cells
BAFF-Rhigh Naive B Cells in Naive B Cells
BAFF-Rhigh B Cells in B Cells
BAFF-Rhigh Switched Memory B Cells in Switched Memory B Cells
CD11c+ CD16− CD56dim NK Cells in CD16− CD56dim NK Cells
BAFF-Rhigh Memory B Cells in Memory B Cells
CD40+ Non-Classical Monocytes in Non−Classical Monocytes
BAFF-Rhigh Transitional B Cells in Transitional B Cells
BAFF-Rhigh DN B Cells in Double Negative B Cells
NK Cells in Live Cells
HLA-DR+ CD38high CD8+ T Cells in CD8+ T Cells
CD86+ Dendritic Cells in Dendritic Cells
CD38+ CD27− CD4+ T Cells in CD27− CD4+ T Cells
CD16+ CD56dim NK Cells in CD16+ NK Cells
CD38+ CD16− CD56dim NK Cells in CD16− CD56dim NK Cells
CD16+ CD56bright NK Cells in NK Cells
CD16+ CD56bright NK Cells in CD16+ NK Cells
CD10+ DN T Cells in Double Negative T Cells
Myeloid DCs in Dendritic Cells
CD86+ mDCs in Myeloid DCs
CD94+ DN T Cells in Double Negative T Cells
CD38+ Non-Classical Monocytes in Non−Classical Monocytes
CD38+ Monocytes in Monocytes
HLA-DR+ NKT Cells in NKT Cells
IgMlow IgD− Naive B Cells in Naive B Cells
IgMlow IgD− Naive B Cells in B Cells
CD38+ CD16− CD56bright NK Cells in CD16− CD56bright NK Cells
HLA-DR+ CD16+ NK Cells in CD16+ NK Cells
CD40+ Intermediate Monocytes in Intermediate Monocytes
CD11c+ Intermediate Monocytes in Intermediate Monocytes
CD16+ CD56dim NK Cells in NK Cells
CD94+ CD16+ NK Cells in CD16+ NK Cells
DN1 B Cells in Double Negative B Cells
CD94+ NK Cells in NK Cells
NKT Cells in Live Cells
CD11c+ Monocytes in Monocytes
HLA-DR+ CD38high CD4+ T Cells in CD4+ T Cells
CD27+ CD16− CD56dim NK Cells in CD16− CD56dim NK Cells
IgM+ IgD− Memory B Cells in Memory B Cells
0.1
0.5
1.0
5.0
10.0
50.0
100.0
OR

**Supplementary Figure 2. Immunological parameters associated with Still's disease.** Patients with Still's disease (n=34) were compared against healthy individuals (n=58) by multivariate logistic regression of immunological parameters. Data from patients with autoinflammation of unknown origin (n=36), FMF (n=35) and Behçet (n=23) are shown as reference data only. **A)** Odds ratio and 95% confidence interval of highly-associated cell population frequency changes in patients with Still's disease in relation to healthy individuals. Estimated by multivariable logistic regression adjusted by sex and age. **B)** Average and 95% confidence interval of 200 times 10 fold cross-validation to evaluate a sufficient number of best cell populations, based on ability to adequately discern between Still's disease and healthy individuals. **C)** Frequency for highly-associated cell populations for Still's disease in relation to healthy individuals. Each dot represents a patient and each colour represents a condition. **D)** Average ROC curve with 95% confidence interval of 10 fold cross-validation for Still's disease in relation to healthy individuals. ROC calculated using multi-variable logistic regression, adjusted by sex and age, considering the 82 cell populations with highest explanatory contribution. Area under ROC curve and confidence interval indicated on graph. **E)** First two PCA components of all cell populations in the dataset. Each dot represents an individual and each colour represents a condition. Histograms show distribution of values in Still's disease and healthy individuals. **F)** First two PCA components of 82 cell populations most highly associated for divergence between Still's disease and healthy individuals. Histograms show distribution of values in Still's disease and healthy individuals. The two arrows show the direction of distinct highly associated cell populations.

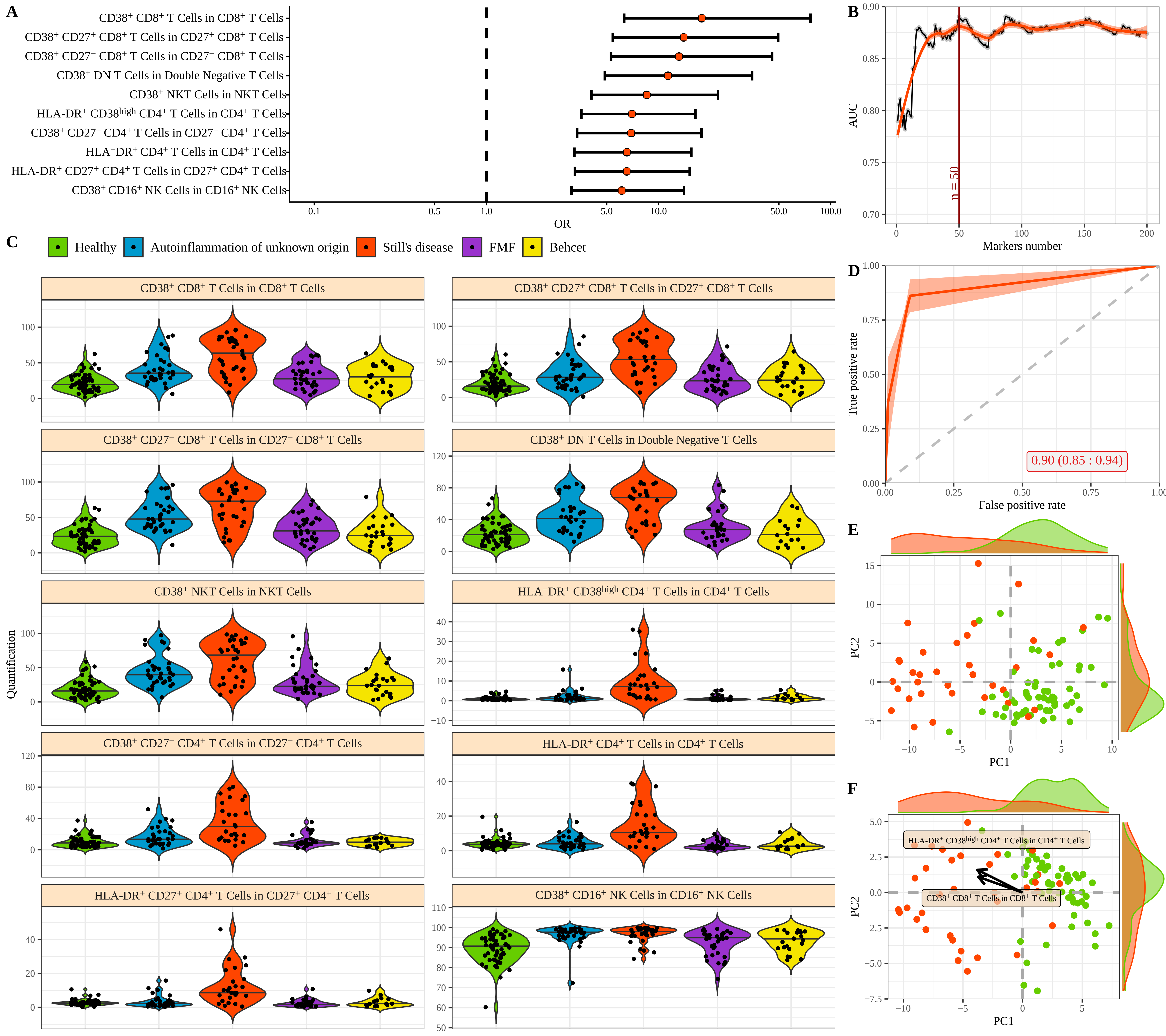
A
CD38+ CD8+ T Cells in CD8+ T Cells
CD38+ CD27+ CD8+ T Cells in CD27+ CD8+ T Cells
CD38+ CD27− CD8+ T Cells in CD27− CD8+ T Cells
CD38+ DN T Cells in Double Negative T Cells
CD38+ NKT Cells in NKT Cells
HLA-DR+ CD38high CD4+ T Cells in CD4+ T Cells
CD38+ CD27− CD4+ T Cells in CD27− CD4+ T Cells
HLA−DR+ CD4+ T Cells in CD4+ T Cells
HLA-DR+ CD27+ CD4+ T Cells in CD27+ CD4+ T Cells
CD38+ CD16+ NK Cells in CD16+ NK Cells
OR
B
AUC
n = 50
Markers number
C
Healthy
Autoinflammation of unknown origin
Still's disease
FMF
Behcet
Quantification
D
True positive rate
False positive rate
0.90 (0.85 : 0.94)
E
PC1
PC2
F
HLA-DR+ CD38high CD4+ T Cells in CD4+ T Cells
CD38+ CD8+ T Cells in CD8+ T Cells

**Supplementary Figure 3. Immunological parameters associated with FMF.** Patients with FMF (n=35) were compared against healthy individuals (n=58) by multivariate logistic regression of immunological parameters. Data from patients with autoinflammation of unknown origin (n=36), Still’s disease (n=34) and Behçet (n=23) are shown as reference data only. **A)** Odds ratio and 95% confidence interval of highly-associated cell population frequency changes in patients with FMF in relation to healthy individuals. Estimated by multivariable logistic regression adjusted by sex and age. **B)** Average and 95% confidence interval of 200 times 10 fold cross-validation to evaluate a sufficient number of best cell populations, based on ability to adequately discern between FMF and healthy individuals. **C)** Frequency for highly-associated cell populations for FMF in relation to healthy individuals. Each dot represents a patient and each colour represents a condition. **D)** Average ROC curve with 95% confidence interval of 10 fold cross-validation for FMF in relation to healthy individuals. ROC calculated using multi-variable logistic regression, adjusted by sex and age, considering the 55 cell populations with highest explanatory contribution. Area under ROC curve and confidence interval indicated on graph. **E)** First two PCA components of all cell populations in the dataset. Each dot represents an individual and each colour represents a condition. Histograms show distribution of values in FMF and healthy individuals. **F)** First two PCA components of **55** cell populations most highly associated for divergence between FMF and healthy individuals. Histograms show distribution of values in FMF and healthy individuals. The two arrows show the direction of distinct highly associated cell populations.

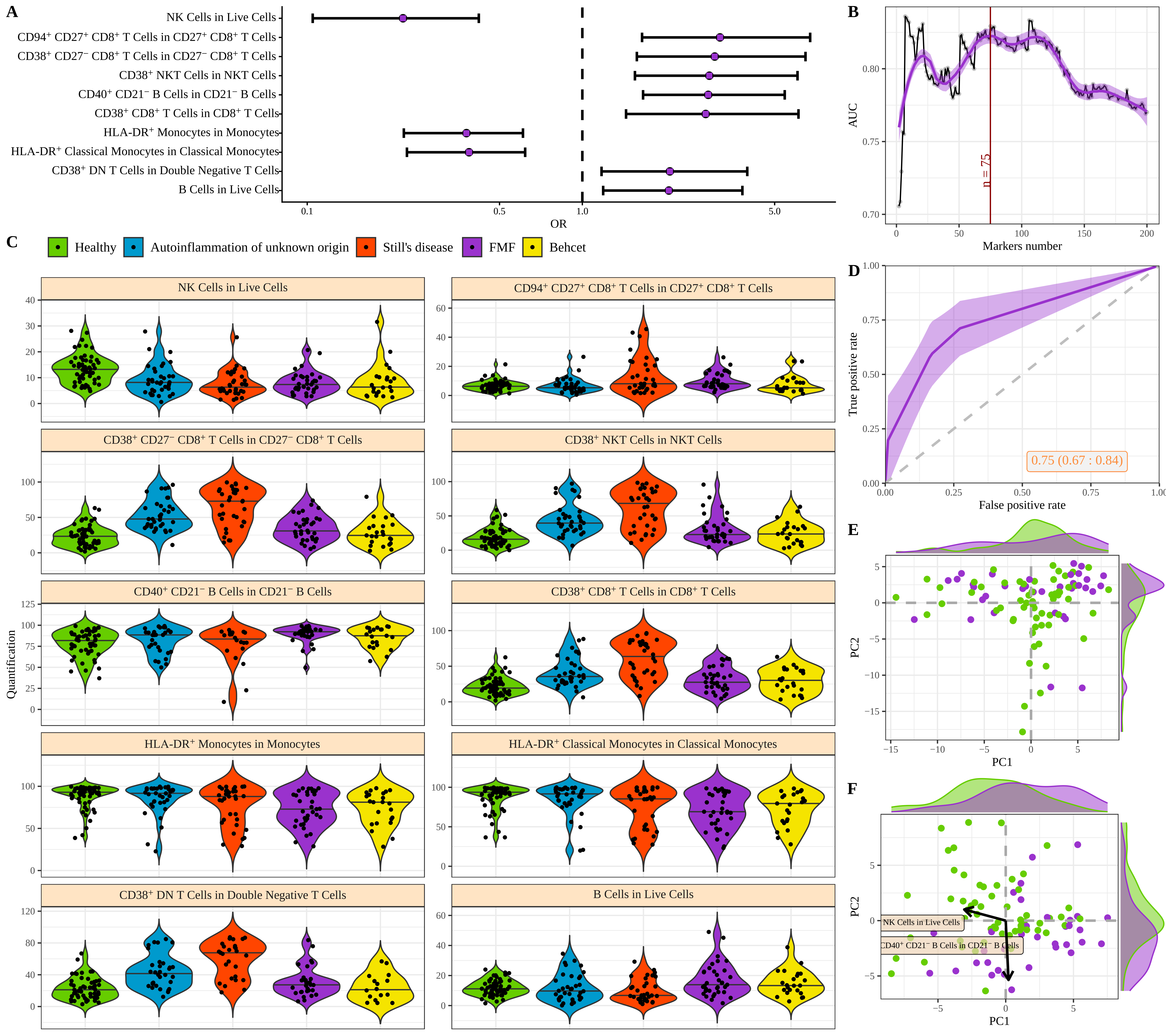
A
NK Cells in Live Cells
CD94+ CD27+ CD8+ T Cells in CD27+ CD8+ T Cells
CD38+ CD27− CD8+ T Cells in CD27− CD8+ T Cells
CD38+ NKT Cells in NKT Cells
CD40+ CD21− B Cells in CD21− B Cells
CD38+ CD8+ T Cells in CD8+ T Cells
HLA-DR+ Monocytes in Monocytes
HLA-DR+ Classical Monocytes in Classical Monocytes
CD38+ DN T Cells in Double Negative T Cells
B Cells in Live Cells
OR
B
AUC
n = 75
Markers number
C
Healthy
Autoinflammation of unknown origin
Still's disease
FMF
Behcet
Quantification
D
True positive rate
False positive rate
0.75 (0.67 : 0.84)
E
PC1
PC2
F
NK Cells in Live Cells
CD40+ CD21− B Cells in CD21− B Cells
PC1
PC2

**Supplementary Figure 4. Immunological parameters associated with Behçet's disease.** Patients with Behçet's disease (n=23) were compared against healthy individuals (n=58) by multivariate logistic regression of immunological parameters. Data from patients with autoinflammation of unknown origin (n=36), Still's disease (n=34) and FMF (n=35) are shown as reference data only. **A)** Odds ratio and 95% confidence interval of highly-associated cell population frequency changes in patients with Behçet in relation to healthy individuals. Estimated by multivariable logistic regression adjusted by sex and age. **B)** Average and 95% confidence interval of 200 times 10 fold cross-validation to evaluate a sufficient number of best cell populations, based on ability to adequately discern between Behçet and healthy individuals. **C)** Frequency for highly-associated cell populations for Behçet in relation to healthy individuals. Each dot represents a patient and each colour represents a condition. **D)** Average ROC curve with 95% confidence interval of 10 fold cross-validation for Behçet in relation to healthy individuals. ROC calculated using multi-variable logistic regression, adjusted by sex and age, considering the 61 cell populations with highest explanatory contribution. Area under ROC curve and confidence interval indicated on graph. **E)** First two PCA components of all cell populations in the dataset. Each dot represents an individual and each colour represents a condition. Histograms show distribution of values in Behçet and healthy individuals. **F)** First two PCA components of 61 cell populations most highly associated for divergence between Behçet and healthy individuals. Histograms show distribution of values in Behçet and healthy individuals. The two arrows show the direction of distinct highly associated cell populations.

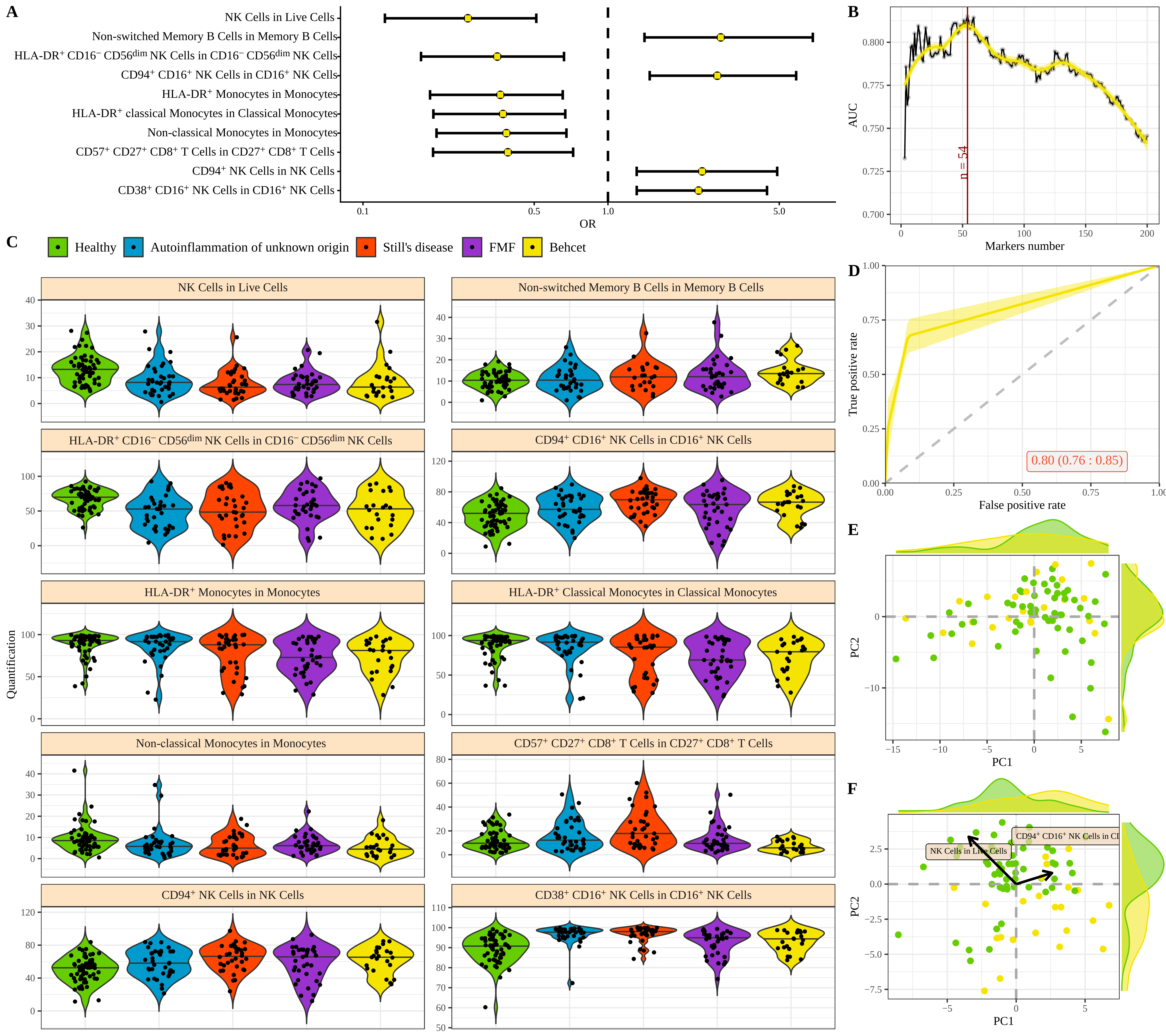
A
NK Cells in Live Cells
Non-switched Memory B Cells in Memory B Cells
HLA-DR+ CD16− CD56dim NK Cells in CD16− CD56dim NK Cells
CD94+ CD16+ NK Cells in CD16+ NK Cells
HLA-DR+ Monocytes in Monocytes
HLA-DR+ classical Monocytes in Classical Monocytes
Non-classical Monocytes in Monocytes
CD57+ CD27+ CD8+ T Cells in CD27+ CD8+ T Cells
CD94+ NK Cells in NK Cells
CD38+ CD16+ NK Cells in CD16+ NK Cells
OR
B
AUC
Markers number
n = 54
C
Healthy
Autoinflammation of unknown origin
Still's disease
FMF
Behcet
Quantification
D
True positive rate
False positive rate
0.80 (0.76 : 0.85)
E
PC1
PC2
F
NK Cells in Live Cells
CD94+ CD16+ NK Cells in CD
PC1
PC2

**Supplementary Figure 5. Key immunological features driving machine learning-led disease identification.** A multi-disease comparison was performed using a Random Forest algorithm to identify immune characteristics with discriminating potential between patients with autoinflammation of unknown origin (n=36), Still's disease (n=34), FMF (n=35) and Behçet's (n=23). Healthy individuals were not used in the model generation, and are shown as reference data only. Model importance for the 50 highest associated cell populations for disease discrimination.

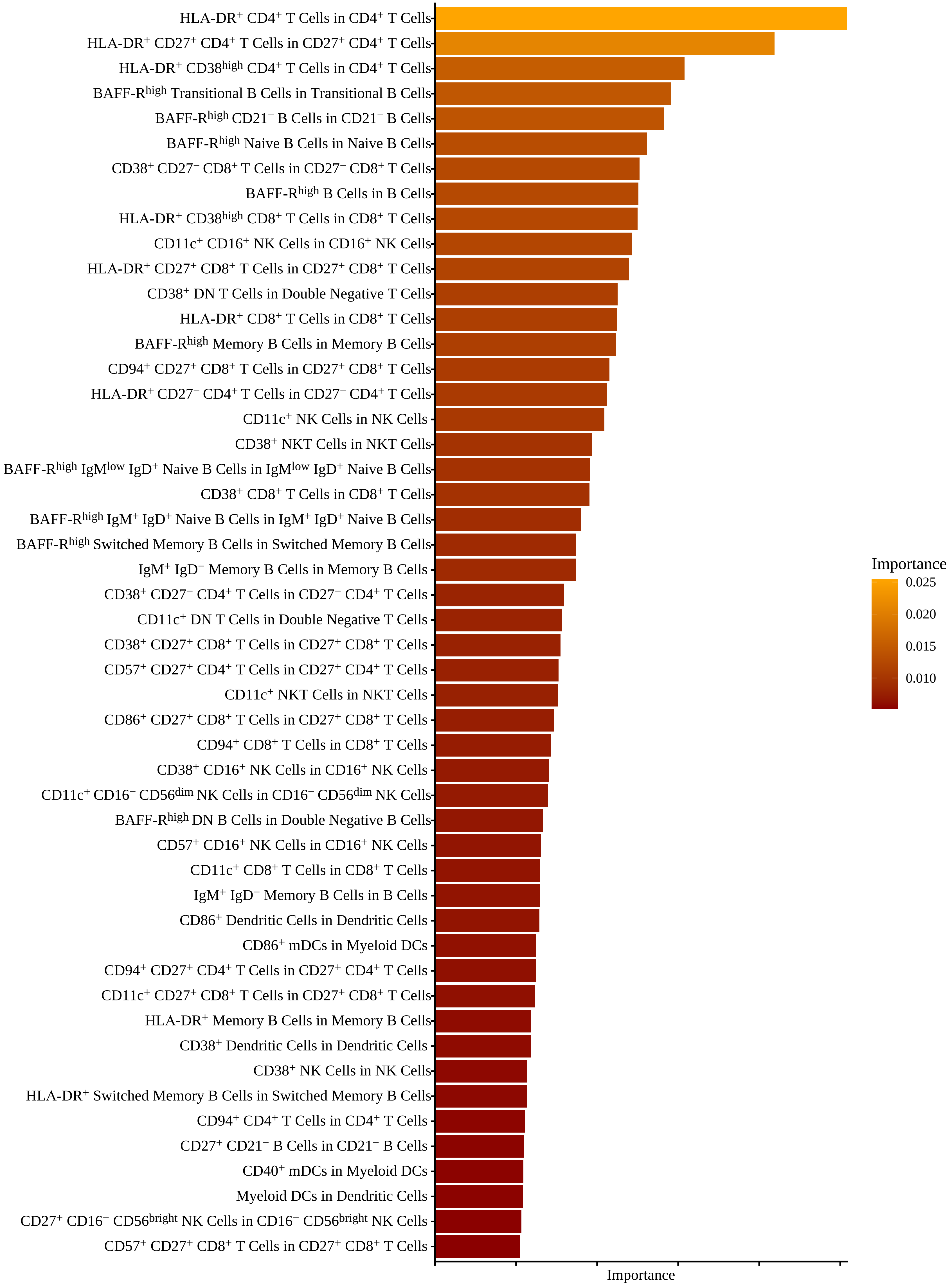
HLA-DR$^{+}$ CD4$^{+}$ T Cells in CD4$^{+}$ T Cells
HLA-DR$^{+}$ CD27$^{+}$ CD4$^{+}$ T Cells in CD27$^{+}$ CD4$^{+}$ T Cells
HLA-DR$^{+}$ CD38$^{high}$ CD4$^{+}$ T Cells in CD4$^{+}$ T Cells
BAFF-R$^{high}$ Transitional B Cells in Transitional B Cells
BAFF-R$^{high}$ CD21$^{-}$ B Cells in CD21$^{-}$ B Cells
BAFF-R$^{high}$ Naive B Cells in Naive B Cells
CD38$^{+}$ CD27$^{-}$ CD8$^{+}$ T Cells in CD27$^{-}$ CD8$^{+}$ T Cells
BAFF-R$^{high}$ B Cells in B Cells
HLA-DR$^{+}$ CD38$^{high}$ CD8$^{+}$ T Cells in CD8$^{+}$ T Cells
CD11c$^{+}$ CD16$^{+}$ NK Cells in CD16$^{+}$ NK Cells
HLA-DR$^{+}$ CD27$^{+}$ CD8$^{+}$ T Cells in CD27$^{+}$ CD8$^{+}$ T Cells
CD38$^{+}$ DN T Cells in Double Negative T Cells
HLA-DR$^{+}$ CD8$^{+}$ T Cells in CD8$^{+}$ T Cells
BAFF-R$^{high}$ Memory B Cells in Memory B Cells
CD94$^{+}$ CD27$^{+}$ CD8$^{+}$ T Cells in CD27$^{+}$ CD8$^{+}$ T Cells
HLA-DR$^{+}$ CD27$^{-}$ CD4$^{+}$ T Cells in CD27$^{-}$ CD4$^{+}$ T Cells
CD11c$^{+}$ NK Cells in NK Cells
CD38$^{+}$ NKT Cells in NKT Cells
BAFF-R$^{high}$ IgM$^{low}$ IgD$^{+}$ Naive B Cells in IgM$^{low}$ IgD$^{+}$ Naive B Cells
CD38$^{+}$ CD8$^{+}$ T Cells in CD8$^{+}$ T Cells
BAFF-R$^{high}$ IgM$^{+}$ IgD$^{+}$ Naive B Cells in IgM$^{+}$ IgD$^{+}$ Naive B Cells
BAFF-R$^{high}$ Switched Memory B Cells in Switched Memory B Cells
IgM$^{+}$ IgD$^{-}$ Memory B Cells in Memory B Cells
CD38$^{+}$ CD27$^{-}$ CD4$^{+}$ T Cells in CD27$^{-}$ CD4$^{+}$ T Cells
CD11c$^{+}$ DN T Cells in Double Negative T Cells
CD38$^{+}$ CD27$^{+}$ CD8$^{+}$ T Cells in CD27$^{+}$ CD8$^{+}$ T Cells
CD57$^{+}$ CD27$^{+}$ CD4$^{+}$ T Cells in CD27$^{+}$ CD4$^{+}$ T Cells
CD11c$^{+}$ NKT Cells in NKT Cells
CD86$^{+}$ CD27$^{+}$ CD8$^{+}$ T Cells in CD27$^{+}$ CD8$^{+}$ T Cells
CD94$^{+}$ CD8$^{+}$ T Cells in CD8$^{+}$ T Cells
CD38$^{+}$ CD16$^{+}$ NK Cells in CD16$^{+}$ NK Cells
CD11c$^{+}$ CD16$^{-}$ CD56$^{dim}$ NK Cells in CD16$^{-}$ CD56$^{dim}$ NK Cells
BAFF-R$^{high}$ DN B Cells in Double Negative B Cells
CD57$^{+}$ CD16$^{+}$ NK Cells in CD16$^{+}$ NK Cells
CD11c$^{+}$ CD8$^{+}$ T Cells in CD8$^{+}$ T Cells
IgM$^{+}$ IgD$^{-}$ Memory B Cells in B Cells
CD86$^{+}$ Dendritic Cells in Dendritic Cells
CD86$^{+}$ mDCs in Myeloid DCs
CD94$^{+}$ CD27$^{+}$ CD4$^{+}$ T Cells in CD27$^{+}$ CD4$^{+}$ T Cells
CD11c$^{+}$ CD27$^{+}$ CD8$^{+}$ T Cells in CD27$^{+}$ CD8$^{+}$ T Cells
HLA-DR$^{+}$ Memory B Cells in Memory B Cells
CD38$^{+}$ Dendritic Cells in Dendritic Cells
CD38$^{+}$ NK Cells in NK Cells
HLA-DR$^{+}$ Switched Memory B Cells in Switched Memory B Cells
CD94$^{+}$ CD4$^{+}$ T Cells in CD4$^{+}$ T Cells
CD27$^{+}$ CD21$^{-}$ B Cells in CD21$^{-}$ B Cells
CD40$^{+}$ mDCs in Myeloid DCs
Myeloid DCs in Dendritic Cells
CD27$^{+}$ CD16$^{-}$ CD56$^{bright}$ NK Cells in CD16$^{-}$ CD56$^{bright}$ NK Cells
CD57$^{+}$ CD27$^{+}$ CD8$^{+}$ T Cells in CD27$^{+}$ CD8$^{+}$ T Cells
Importance
Importance
0.025
0.020
0.015
0.010

**Supplementary Figure 6. Plasma proteomic parameters with high association with autoinflammation of unknown**. Odds ratio and 95% confidence interval of 50 highly-associated plasma proteomic parameters in patients with inflammation of unknown origin in relation to healthy individuals. Estimated by multivariable logistic regression adjusted by sex and age

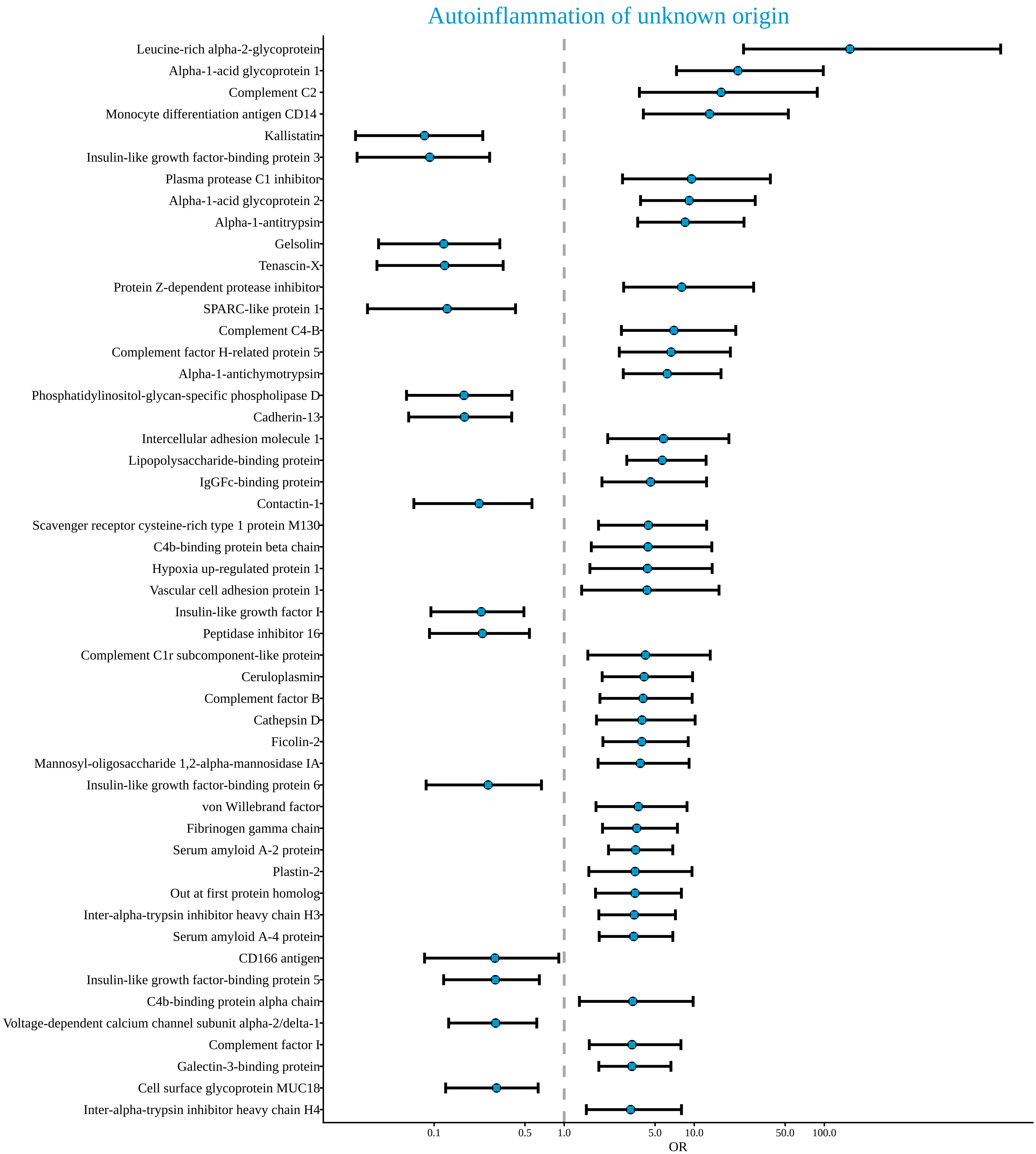
Autoinflammation of unknown origin
Leucine-rich alpha-2-glycoprotein
Alpha-1-acid glycoprotein 1
Complement C2
Monocyte differentiation antigen CD14
Kallistatin
Insulin-like growth factor-binding protein 3
Plasma protease C1 inhibitor
Alpha-1-acid glycoprotein 2
Alpha-1-antitrypsin
Gelsolin
Tenascin-X
Protein Z-dependent protease inhibitor
SPARC-like protein 1
Complement C4-B
Complement factor H-related protein 5
Alpha-1-antichymotrypsin
Phosphatidylinositol-glycan-specific phospholipase D
Cadherin-13
Intercellular adhesion molecule 1
Lipopolysaccharide-binding protein
IgGFc-binding protein
Contactin-1
Scavenger receptor cysteine-rich type 1 protein M130
C4b-binding protein beta chain
Hypoxia up-regulated protein 1
Vascular cell adhesion protein 1
Insulin-like growth factor I
Peptidase inhibitor 16
Complement C1r subcomponent-like protein
Ceruloplasmin
Complement factor B
Cathepsin D
Ficolin-2
Mannosyl-oligosaccharide 1,2-alpha-mannosidase IA
Insulin-like growth factor-binding protein 6
von Willebrand factor
Fibrinogen gamma chain
Serum amyloid A-2 protein
Plastin-2
Out at first protein homolog
Inter-alpha-trypsin inhibitor heavy chain H3
Serum amyloid A-4 protein
CD166 antigen
Insulin-like growth factor-binding protein 5
C4b-binding protein alpha chain
Voltage-dependent calcium channel subunit alpha-2/delta-1
Complement factor I
Galectin-3-binding protein
Cell surface glycoprotein MUC18
Inter-alpha-trypsin inhibitor heavy chain H4
0.1
0.5
1.0
5.0
10.0
50.0
100.0
OR

**Supplementary Figure 7. Plasma proteomic parameters associated with Still's disease.** Patients with Still's disease (n=34) were compared against healthy individuals (n=58) by multivariate logistic regression of immunological parameters. Data from patients with autoinflammation of unknown origin (n=36), FMF (n=35) and Behçet (n=23) are shown as reference data only. **A)** Odds ratio and 95% confidence interval of highly-associated plasma protein changes in patients with Still's disease in relation to healthy individuals. Estimated by multivariable logistic regression adjusted by sex and age. **B)** Average and 95% confidence interval of 200 times 10 fold cross-validation to evaluate a sufficient number of plasma protein changes, based on ability to adequately discern between Still's disease and healthy individuals. **C)** Frequency for highly-associated plasma protein changes for Still's disease in relation to healthy individuals. Each dot represents a patient and each colour represents a condition. **D)** Average ROC curve with 95% confidence interval of 10 fold cross-validation for Still's disease in relation to healthy individuals. ROC calculated using multi-variable logistic regression, adjusted by sex and age, considering the 53 plasma proteins with highest explanatory contribution. Area under ROC curve and confidence interval indicated on graph. **E)** First two PCA components of all plasma proteins in the dataset. Each dot represents an individual and each colour represents a condition. Histograms show distribution of values in Still's disease and healthy individuals. **F)** First two PCA components of 53 plasma proteins most highly associated for divergence between Still's disease and healthy individuals. Histograms show distribution of values in Still's disease and healthy individuals. The two arrows show the direction of distinct highly associated plasma proteins.

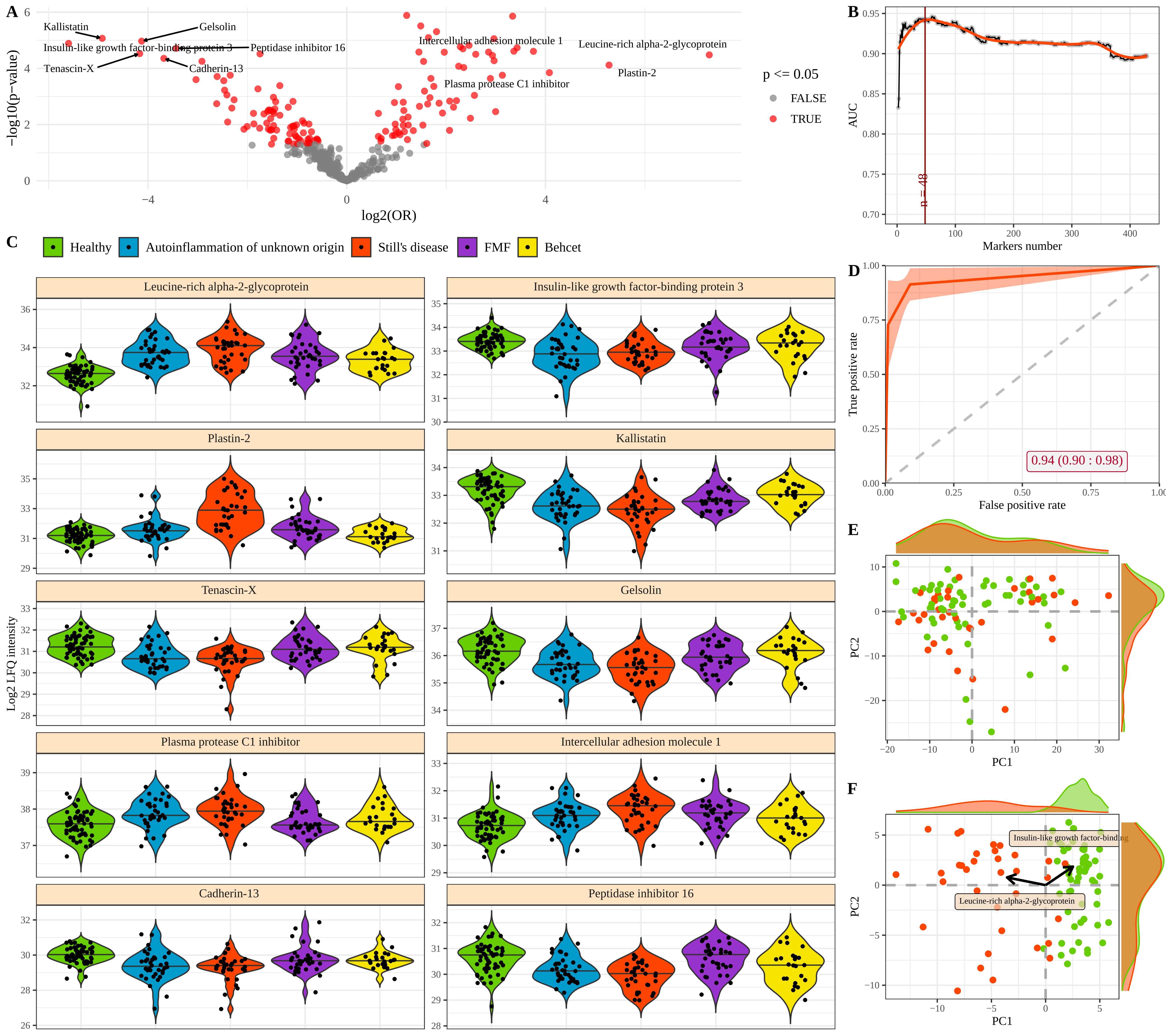
A
Kallistatin
Gelsolin
Insulin-like growth factor-binding protein 3
Peptidase inhibitor 16
Tenascin-X
Cadherin-13
Intercellular adhesion molecule 1
Leucine-rich alpha-2-glycoprotein
Plastin-2
Plasma protease C1 inhibitor
−log10(p−value)
log2(OR)
p <= 0.05
FALSE
TRUE
B
AUC
Markers number
n = 48
C
Healthy
Autoinflammation of unknown origin
Still's disease
FMF
Behcet
Leucine-rich alpha-2-glycoprotein
Insulin-like growth factor-binding protein 3
Plastin-2
Kallistatin
Tenascin-X
Gelsolin
Plasma protease C1 inhibitor
Intercellular adhesion molecule 1
Cadherin-13
Peptidase inhibitor 16
Log2 LFQ intensity
D
True positive rate
False positive rate
0.94 (0.90 : 0.98)
E
PC1
PC2
F
Insulin-like growth factor-binding
Leucine-rich alpha-2-glycoprotein
PC1
PC2

**Supplementary Figure 8. Plasma protein changes associated with FMF.** Patients with FMF (n=35) were compared against healthy individuals (n=58) by multivariate logistic regression of plasma protein parameters. Data from patients with autoinflammation of unknown origin (n=36), Still's disease (n=34) and Behçet (n=23) are shown as reference data only. **A)** Odds ratio and 95% confidence interval of highly-associated plasma protein abundance changes in patients with FMF in relation to healthy individuals. Estimated by multivariable logistic regression adjusted by sex and age. **B)** Average and 95% confidence interval of 200 times 10 fold cross-validation to evaluate a sufficient number of best plasma proteins, based on ability to adequately discern between FMF and healthy individuals. **C)** Frequency for highly-associated plasma proteins for FMF in relation to healthy individuals. Each dot represents a patient and each colour represents a condition. **D)** Average ROC curve with 95% confidence interval of 10 fold cross-validation for FMF in relation to healthy individuals. ROC calculated using multi-variable logistic regression, adjusted by sex and age, considering the 45 plasma proteins with highest explanatory contribution. Area under ROC curve and confidence interval indicated on graph. **E)** First two PCA components of all plasma proteins in the dataset. Each dot represents an individual and each colour represents a condition. Histograms show distribution of values in FMF and healthy individuals. **F)** First two PCA components of 45 plasma proteins most highly associated for divergence between FMF and healthy individuals. Histograms show distribution of values in FMF and healthy individuals. The two arrows show the direction of distinct highly associated plasma proteins.

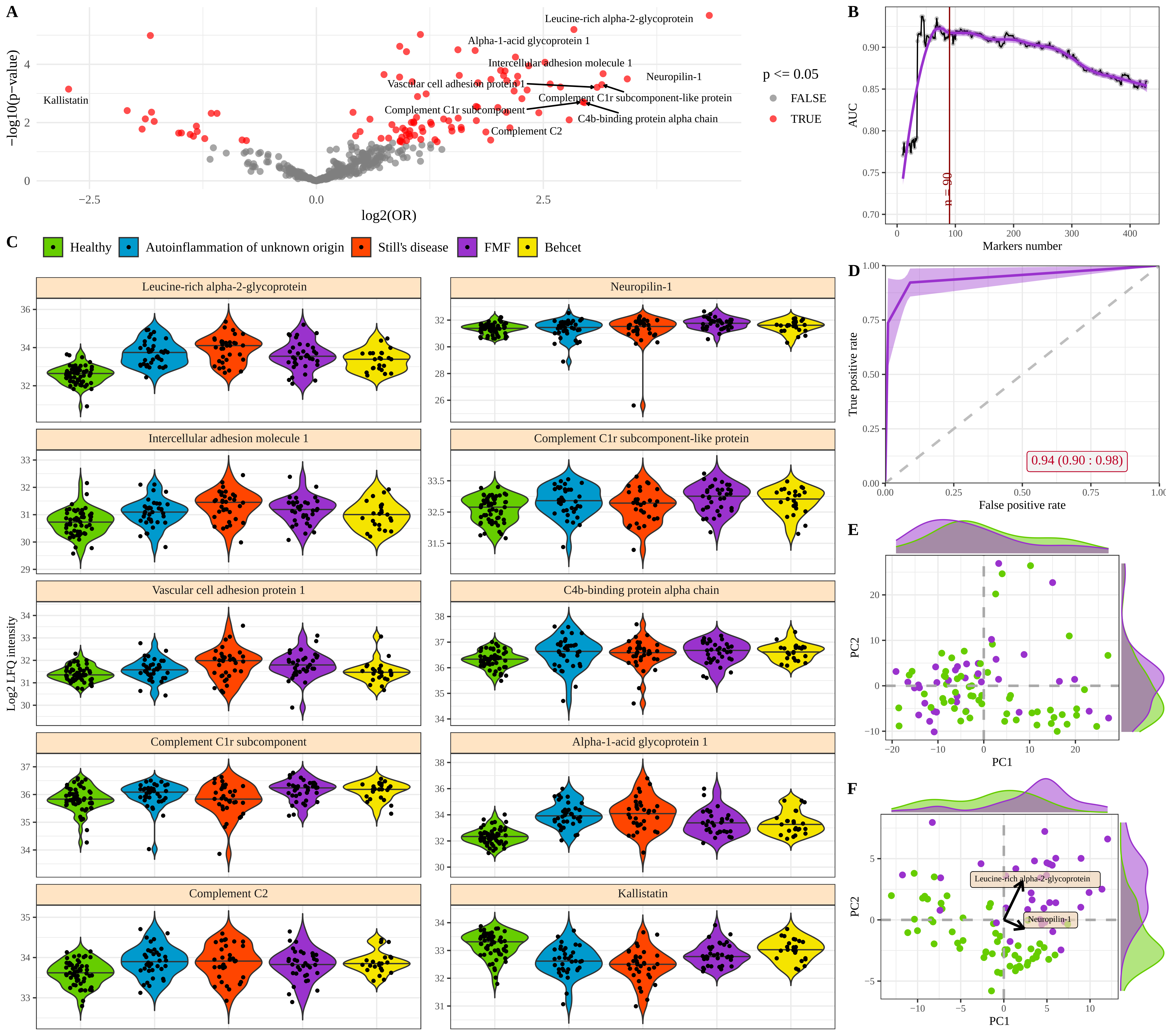
A
B
C
D
E
F
Leucine-rich alpha-2-glycoprotein
Alpha-1-acid glycoprotein 1
Intercellular adhesion molecule 1
Neuropilin-1
Vascular cell adhesion protein 1
Complement C1r subcomponent-like protein
Complement C1r subcomponent
C4b-binding protein alpha chain
Complement C2
Kallistatin
−log10(p−value)
log2(OR)
p <= 0.05
FALSE
TRUE
AUC
Markers number
n = 90
Healthy
Autoinflammation of unknown origin
Still's disease
FMF
Behcet
Log2 LFQ intensity
True positive rate
False positive rate
0.94 (0.90 : 0.98)
PC1
PC2
Neuropilin-1

**Supplementary Figure 9. Plasma protein changes associated with Behçet's disease.** Patients with Behçet's disease (n=23) were compared against healthy individuals (n=58) by multivariate logistic regression of plasma proteomic parameters. Data from patients with autoinflammation of unknown origin (n=36), Still's disease (n=34) and FMF (n=35) are shown as reference data only. **A)** Odds ratio and 95% confidence interval of highly-associated plasma protein abundance changes in patients with Behçet in relation to healthy individuals. Estimated by multivariable logistic regression adjusted by sex and age. **B)** Average and 95% confidence interval of 200 times 10 fold cross-validation to evaluate a sufficient number of best plasma proteins, based on ability to adequately discern between Behçet and healthy individuals. **C)** Frequency for highly-associated plasma proteins for Behçet in relation to healthy individuals. Each dot represents a patient and each colour represents a condition. **D)** Average ROC curve with 95% confidence interval of 10 fold cross-validation for Behçet in relation to healthy individuals. ROC calculated using multi-variable logistic regression, adjusted by sex and age, considering the 112 plasma proteins with highest explanatory contribution. Area under ROC curve and confidence interval indicated on graph. **E)** First two PCA components of all plasma proteins in the dataset. Each dot represents an individual and each colour represents a condition. Histograms show distribution of values in Behçet and healthy individuals. **F)** First two PCA components of 112 plasma proteins most highly associated for divergence between Behçet and healthy individuals. Histograms show distribution of values in Behçet and healthy individuals. The two arrows show the direction of distinct highly associated plasma proteins.

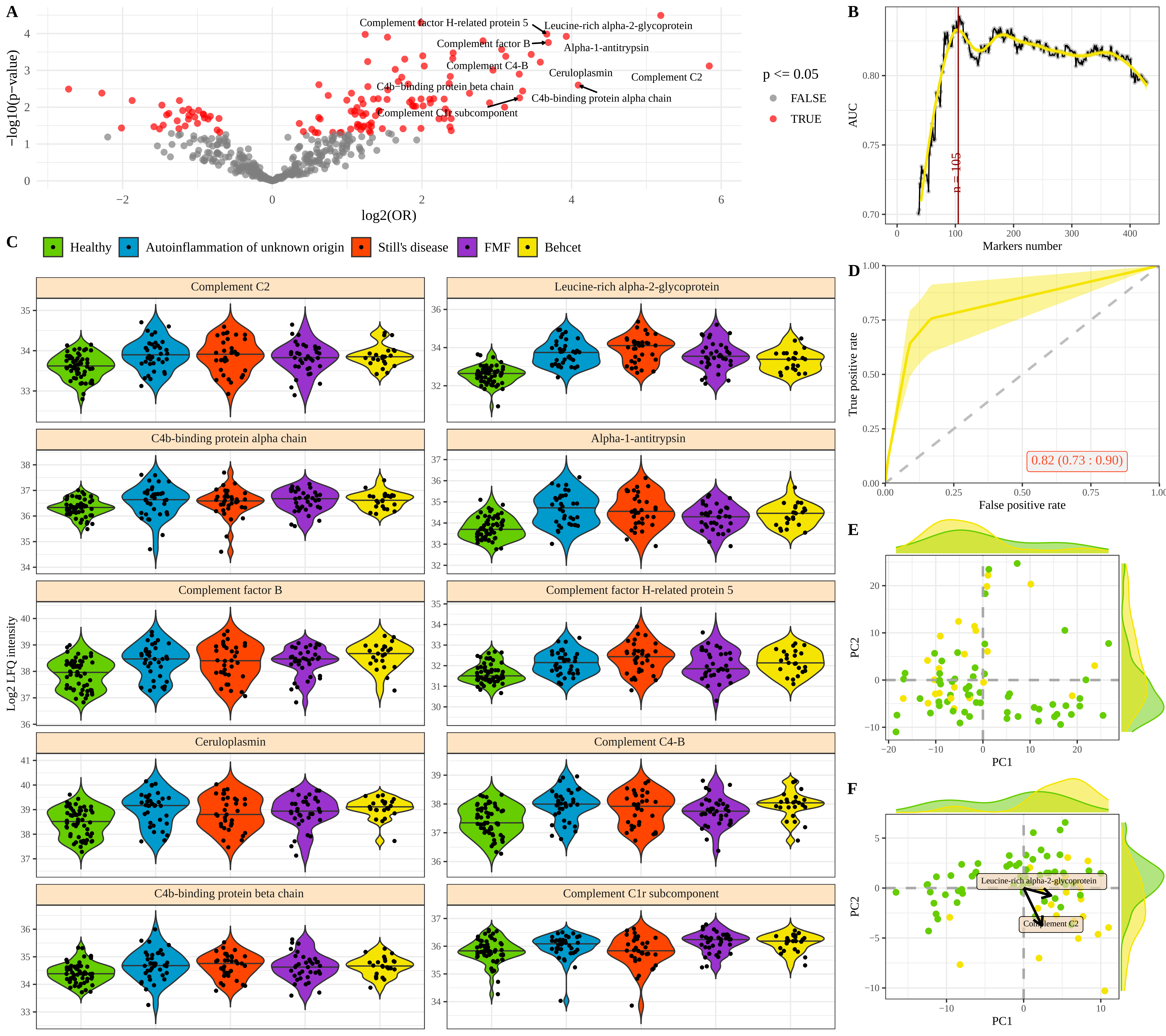
A
Complement factor H-related protein 5
Leucine-rich alpha-2-glycoprotein
Complement factor B
Alpha-1-antitrypsin
Complement C4-B
Ceruloplasmin
Complement C2
C4b-binding protein beta chain
C4b-binding protein alpha chain
Complement C1r subcomponent
−log10(p−value)
log2(OR)
p <= 0.05
FALSE
TRUE
B
AUC
Markers number
n = 105
C
Healthy
Autoinflammation of unknown origin
Still's disease
FMF
Behcet
Log2 LFQ intensity
D
True positive rate
False positive rate
0.82 (0.73 : 0.90)
E
PC1
PC2
F
PC1
PC2
Leucine-rich alpha-2-glycoprotein
Complement C2

**Supplementary Figure 10. Key proteomic features driving machine learning-led disease identification.** A multi-disease comparison was performed using a Random Forest algorithm to identify plasma proteomic characteristics with discriminating potential between patients with autoinflammation of unknown origin (n=36), Still's disease (n=34), FMF (n=35) and Behçet's (n=23). Healthy individuals were not used in the model generation, and are shown as reference data only. Model importance for the 50 highest associated plasma protein changes for disease discrimination.

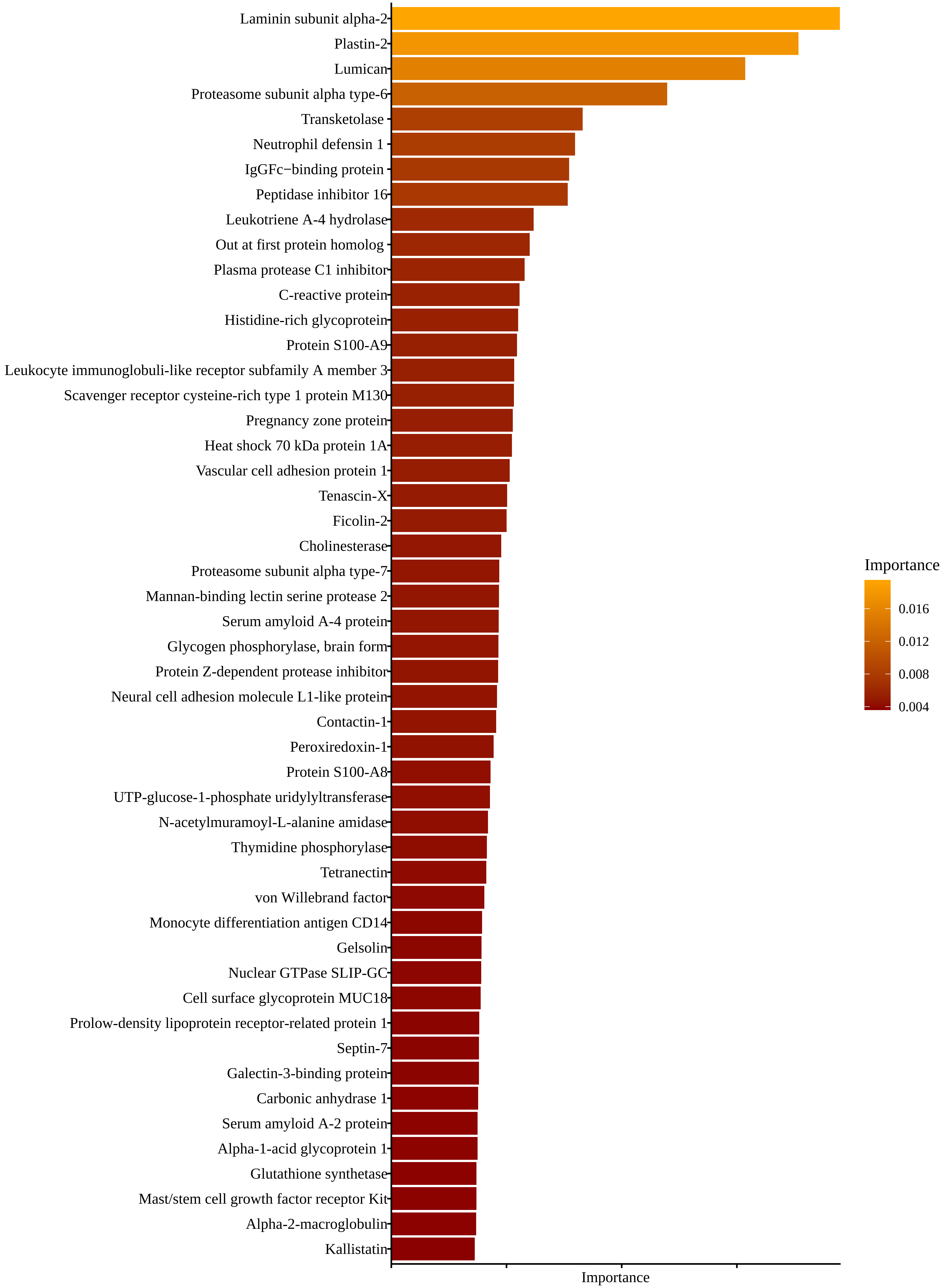
Laminin subunit alpha-2
Plastin-2
Lumican
Proteasome subunit alpha type-6
Transketolase
Neutrophil defensin 1
IgGFc−binding protein
Peptidase inhibitor 16
Leukotriene A-4 hydrolase
Out at first protein homolog
Plasma protease C1 inhibitor
C-reactive protein
Histidine-rich glycoprotein
Protein S100-A9
Leukocyte immunoglobuli-like receptor subfamily A member 3
Scavenger receptor cysteine-rich type 1 protein M130
Pregnancy zone protein
Heat shock 70 kDa protein 1A
Vascular cell adhesion protein 1
Tenascin-X
Ficolin-2
Cholinesterase
Proteasome subunit alpha type-7
Mannan-binding lectin serine protease 2
Serum amyloid A-4 protein
Glycogen phosphorylase, brain form
Protein Z-dependent protease inhibitor
Neural cell adhesion molecule L1-like protein
Contactin-1
Peroxiredoxin-1
Protein S100-A8
UTP-glucose-1-phosphate uridylyltransferase
N-acetylmuramoyl-L-alanine amidase
Thymidine phosphorylase
Tetranectin
von Willebrand factor
Monocyte differentiation antigen CD14
Gelsolin
Nuclear GTPase SLIP-GC
Cell surface glycoprotein MUC18
Prolow-density lipoprotein receptor-related protein 1
Septin-7
Galectin-3-binding protein
Carbonic anhydrase 1
Serum amyloid A-2 protein
Alpha-1-acid glycoprotein 1
Glutathione synthetase
Mast/stem cell growth factor receptor Kit
Alpha-2-macroglobulin
Kallistatin
Importance
Importance
0.016
0.012
0.008
0.004
